# Longitudinal analysis of blood markers reveals progressive loss of resilience and predicts ultimate human lifespan limit

**DOI:** 10.1101/618876

**Authors:** Timothy V. Pyrkov, Konstantin Avchaciov, Andrei E. Tarkhov, Leonid I. Menshikov, Andrei V. Gudkov, Peter O. Fedichev

## Abstract

We investigated the dynamic properties of the organism state fluctuations along individual aging trajectories in a large longitudinal database of CBC measurements from a consumer diagnostics laboratory. To simplify the analysis, we used a log-linear mortality estimate from the CBC variables as a single quantitative measure of aging process, henceforth referred to as dynamic organism state index (DOSI). We observed, that the age-dependent population DOSI distribution broadening could be explained by a progressive loss of physiological resilience measured by the DOSI auto-correlation time. Extrapolation of this trend suggested that DOSI recovery time and variance would simultaneously diverge at a critical point of 120 − 150 years of age corresponding to a complete loss of resilience. The observation was immediately confirmed by the independent analysis of correlation properties of intraday physical activity levels fluctuations collected by wearable devices. We conclude that the criticality resulting in the end of life is an intrinsic biological property of an organism that is independent of stress factors and signifies a fundamental or absolute limit of human lifespan.

## INTRODUCTION

Aging is manifested as a progressive functional decline leading to exponentially increasing prevalence [1, 2] and incidence of chronic age-related diseases (e.g., cancers, diabetes, cardiovascular diseases, etc. [3–5]) and disease-specific mortality [6]. Much of our current understanding of the relationship between aging and changes in physiological variables over an organism’s lifespan originates from large cross-sectional studies [7–9] and led to development of increasingly reliable “biological clocks” or “biological age” estimations reflecting age-related variations in blood markers [10], DNA methylation (DNAm) states [11, 12] or patterns of locomotor activity [13–15] (see [16] for a review of biological age predictors). All-cause mortality in humans [17, 18] and the incidence of chronic agerelated diseases increase exponentially and double every eight years [3]. Typically, however, the physiological indices and the derived quantities such as biological age predictions change from the levels observed in the young organism at a much lower pace than it could be expected from the Gompertzian mortality acceleration.

Most important factors that are strongly associated with age are also known as the hallmarks of aging [19] and may be, at least in principle, modified pharmacologically. In addition to that, by analogy to resilience in ecological systems, the dynamic properties such as physiological resilience measured as the recovery rate from the organism state perturbations [20, 21] were also associated with mortality [22] and thus may serve as an early warning sign of impending health outcomes [23, 24]. Hence, a better quantitative understanding of the intricate relationship between the slow physiological state dynamics, resilience, and the exponential morbidity and mortality acceleration is required to allow the rational design, development, and clinical validation of effective anti-aging interventions.

We addressed these theoretical and practical issues by a systematic investigation of aging, organism state fluctuations, and gradual loss of resilience in a dataset involving multiple Complete Blood Counts (CBC) measured over short periods of time (a few months) from the same person along the individual aging trajectory. To simplify the matters, we followed [25, 26] and described the organism state by means of a single variable, henceforth referred to as the dynamic organism state index (DOSI) in the form of the log-transformed proportional all-cause mortality model predictor. First, we observed that early in life the DOSI dynamics quantitatively follows the universal ontogenetic growth trajectory from [27]. Once the growth phase is completed, the indicator demonstrated all the expected biological age properties, such as association with age, multiple morbidity, unhealthy lifestyles, mortality and future incidence of chronic diseases.

Late in life, the dynamics of the organism state captured by DOSI along the individual aging trajectories is consistent with that of a stochastic process (random walk) on top of the slow aging drift. The increase in the DOSI variability is approximately linear with age and can be explained by the rise of the organism state recovery time. The latter is thus an independent biomarker of aging and a characteristic of resilience. Our analysis shows that the auto-correlation time of DOSI fluctuations grows (and hence the recovery rate decreases) with age from about 2 weeks to over 8 weeks for cohorts aging from 40 to 90 years. The divergence of the recovery time at advanced ages appeared to be an organism-level phenomenon. This was independently confirmed by the investigation of the variance and the autocorrelation properties of physical activity levels from another longitudinal dataset of intraday step-counts measured by wearable devices. We put forward arguments suggesting that such behavior is typical for complex systems near a bifurcation (disintegration) point and thus the progressive loss of resilience with age may be a dynamic origin of the Gompertz law. Finally, we noted, by extrapolation, that the recovery time would diverge and hence the resilience would be ultimately lost at the critical point at the age in the range of 120 − 150 years, thus indicating the absolute limit of human lifespan.

## RESULTS

### Quantification of aging and development

Complete blood counts (CBC) measurements are most frequently included in standard blood tests and thus comprise a large common subset of physiological indices reported across UKB (471473 subjects, age range 39 − 73 y.o.) and NHANES datasets (72925 subjects, age range 1 − 85 y.o., see Table S1 for the description of the data fields). To understand the character of age-related evolution of the organism state we employed a convenient dimensionality-reduction technique, the Principal Component Analysis (PCA). The coordinates of each point in Fig. 1A is obtained by averaging the first three Principal Component scores of PCA-transformed CBC variables in age-matched cohorts in NHANES dataset. The average points follow a well-defined trajectory or a flow in the multivariate configuration space spanned by the physiological variables and clearly correspond to various stages of the organism development and aging.

**FIG. 1:**
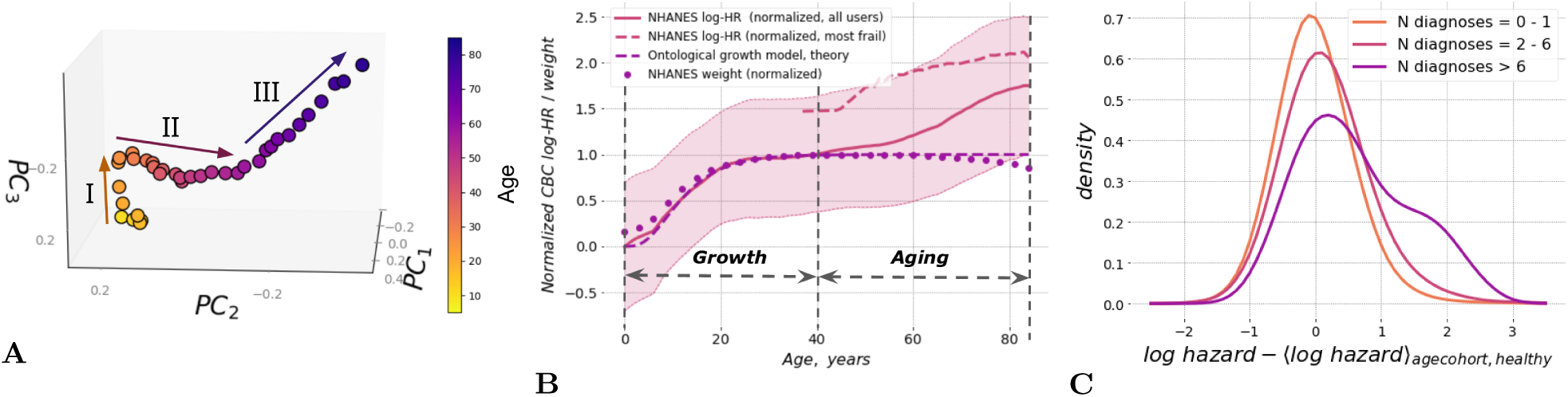
**A**. The graphical representation of the PCA for 5 − 85 year old NHANES participants follows an age-cohort averaged aging trajectory. Centers of each sequential age cohort are plotted in first three PCs. Three approximately linear segments are clearly seen in aging trajectory, corresponding to I) age<35; II) age 35-65; III) age>65. **B**. Dynamic organism state indicator (DOSI) mean values (solid line) and variance (shaded area) are plotted relative to age for all participants of NHANES study. The average line demonstrates nearly linear growth after age of 40. In younger ages the dependence of age is different and consistent with the universal curve suggested by the general model for ontogenetic growth [27]. To illustrate the general character of this early-life dependence we superimposed it with the curve of mean weight in age cohorts of the same population (dotted line). All values are plotted in normalized from as in [27]. The average DOSI of the “most frail” (CMI *>* 0.6) individuals is shown with the dashed line. **C**. Distributions of sex- and age-adjusted DOSI in cohorts of NHANES participants in different morbidity categories relative to the DOSI mean in cohorts of “non-frail” (1 or no diagnoses, CMI *<* 0.1) individuals. Note that the distribution function in the “most frail” group (more than 6 diagnoses, CMI *>* 0.6) exhibited the largest shift and a profound deviation from the symmetric form.

Qualitatively, we differentiated three distinctive segments of the aging trajectory, corresponding to (I) early adulthood (16 − 35 y.o.); (II) middle ages (35 − 65 y.o.); and (III) older ages (older than 65 y.o.). Inside each of the segments, the trajectory was approximately linear. This suggests that over long periods of time (age), CBC variations other than noise could be described by the dynamics of a single dynamic variable (degree of freedom) tracking the distance travelled along the aging trajectory and henceforth referred to as the dynamic organism state indicator (DOSI).

Morbidity and mortality rates increase exponentially with age and a log-linear risk predictor model is a good starting point for characterization of the functional state of an organism and quantification of the aging process [15, 25]. Accordingly, we employed Cox proportional hazards model [28] and trained it using the death register of the NHANES study using log-transformed CBC measurements and sex variable (but not age) as covariates. Altogether, the training subset comprised 23814 participants aged 40 y.o. and older with 3799 death events observed in the follow-up by year 2015. The mortality risk model yielded a single value of log-hazards ratio for every subject and increased in full age range of NHANES participants (Figure 1B). As we will see below, it was a useful and dynamic measure of the organism state henceforth identified with DOSI.

Early in life the dynamics of the organism state has, of course, nothing to do with the late-life increase of mortality rate (i.e., aging), but is rather associated with ontogenetic growth. Accordingly, we checked that the organism state measured by DOSI follows closely the theoretical trajectory of the body mass adopted from [27]:

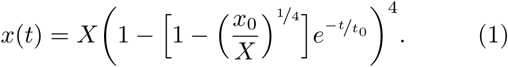

Here *x* is the body mass, or in the linear regime any quantity such as DOSI depending on the body mass, *t* is the age, *t*_0_ is the characteristic time scale associated with the development, and *x*_0_ and *X* are the asymptotic levels of same property at birth and in the fully grown state, respectively. The dots and the dashed lines in Fig. 1B represents age-cohorts averaged body mass trajectory and the best fit of the age-cohort averaged DOSI levels by Eq. (1) for the same NHANES participants. The approximation works remarkably well up until the age of about 40. The characteristic time scale from the fit, *t*_0_ = 6.8 years, coincides almost exactly with the best fit value of 6.3 years obtained from the fit of body mass trajectory.

As the body size increases, the metabolic output per unit mass slows down and the organism reaches a steady state corresponding to the fully grown organism. The inspection of Fig. 1B shows, however, that the equilibrium solution of the organism growth problem appears to be unstable in the long run and the organism state dynamics measured by DOSI exhibits deviations from the stationary solution beyond the age of approximately 40 years old.

To separate the effects of chronic diseases from disease-free aging, we followed [29] and characterized the health status of each study participant based the number of health conditions diagnosed for an individual normalized to the total number of conditions included in the analysis to yield the “compound morbidity index” (CMI) with values ranging from zero to one. The list of health conditions common to the NHANES and UKB studies that were used for CMI determination is given in Table S2 and described in Supplementary Information.

Multiple morbidity manifests itself as elevated DOSI levels. This can be readily seen from the difference between the solid and dashed lines in Fig. 1B, which represent the DOSI means in the cohorts of healthy (“non-frail”, CMI *<* 0.1) and “most frail” (CMI *>* 0.6) NHANES participants, respectively. In groups stratified by increasing number of health condition diagnoses, the normalized distribution of DOSI values (after adjustment by the respective mean levels in age- and sex-matched cohorts of healthy subjects) exhibited a progressive shift and increased variability (see Figs. 1C and S1B for NHANES and UKB, respectively).

For both NHANES and UKB, the largest shift was observed in the “most frail” (CMI *>* 0.6) population. The increasingly heavy tail at the high end of the DOSI distribution in this group is characteristic of an admixture of a distinct group of individuals occupying the adjacent region in the configuration space corresponding to the largest possible DOSI levels. Therefore, DOSI displacement from zero mean (after proper adjustments for age and sex) was expected to reflect the fraction of “most frail” individuals in a cohort of any given age. This was confirmed to be true using the NHANES dataset (Fig. 2A; r = 0.91, p = 2.6 *×* 10^−11^).

**FIG. 2:**
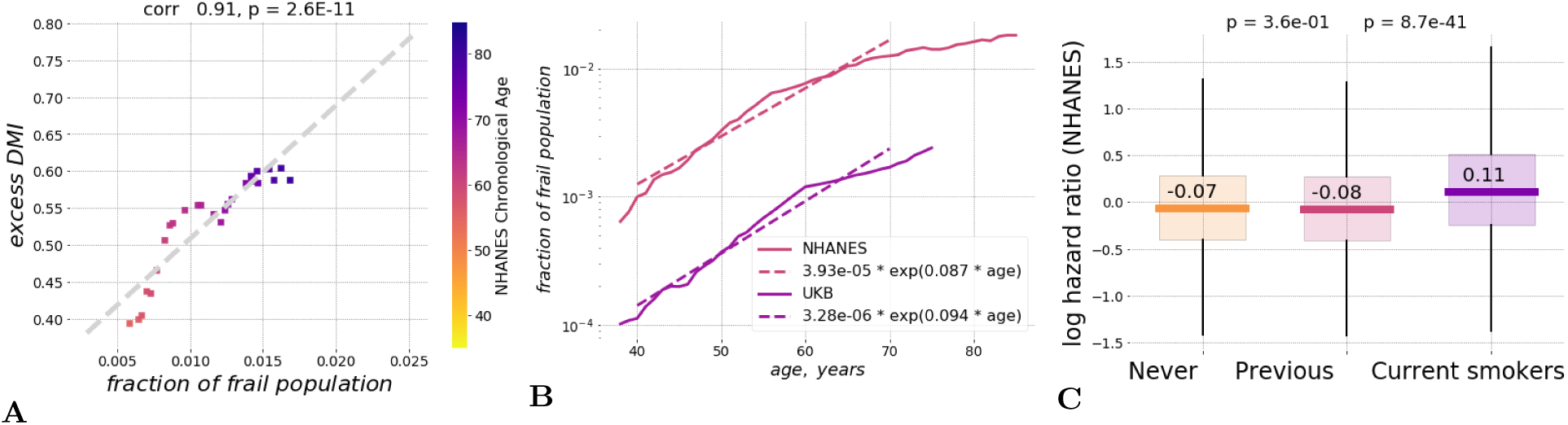
**A**. Fraction of frail persons is strongly correlated with the excess DOSI levels, that is the difference between the DOSI of an individual and its average and the sex- and age-matched cohort in the “non-frail” population in NHANES. **B**. Exponential fit showed that until the age of 70 y.o. the fraction of the “most frail” individuals in the population grows approximately exponentially with age with the doubling rate constants of 0.08 and 0.10 per year in the NHANES and the UKB cohorts, respectively. **C**. Distribution of log hazards ratio in age- and sex-matched cohorts of NHANES participants who never smoked, smoked previously but quit prior to the time of study participation, or were current smokers at the time of the study. The DOSI level is elevated for current smokers, while it is almost indistinguishable between never-smokers and those who quit smoking (p > 0.05).

The fraction of “most frail” subjects still alive increased exponentially at every given age until the age corresponding to the end of healthspan was reached. The characteristic doubling rate constants for the “most frail” population fractions were 0.087 and 0.094 per year in the NHANES and the UKB cohorts, respectively, in comfortable agreement with the accepted Gompertz mortality doubling rate of 0.085 per year [30], see Fig. 2B.

We note that the prevalence of diseases in the NHANES cohort is consistently higher than that in the UKB population, although the average lifespan is comparable in the two countries. This may be a consequence of the enrollment bias in the UKB: life tables analysis in [31] suggests the UKB subjects appear to outlive typical UK residents.

### Dynamic organism state index (DOSI) and health risks

In the most healthy subjects, i.e. those with no diagnosed diseases at the time of assessment, the DOSI predicted the future incidence of chronic age-related diseases observed during 10-year follow-up in the UB study (Table S2). There was no relevant information available in NHANES. We tested this association using a series of Cox proportional hazard models trained to predict the age at the onset/diagnosis of specific diseases. We observed that the morbidity hazard ratios associated with the DOSI relative to its mean in age- and sex-matched cohorts were statistically significant predictors for at least the most prevalent health conditions (those with more than 3000 occurrences in the UKB population). The effect size (HR *≈* 1.03 − 1.07) was the same regardless of whether a disease was diagnosed first in a given individual or followed any number of other diseases. Only emphysema and heart failure which are known to be strongly associated with increased neutrophil counts [32, 33] demonstrated particularly high associations. Therefore, we conclude that the DOSI is a characteristic of overall health status that is universally associated with the risks of developing the most prevalent diseases and, therefore, with the end of healthspan as indicated by the onset of the first morbidity (HR *≈* 1.05 for the “First morbidity” entry in Table S2).

In the most healthy “non-frail” individuals with life-shortening lifestyles/behaviors, such as smoking, the DOSI was also elevated, indicating a higher level of risks of future diseases and death (Fig. 2C). Notably and in agreement with the dynamic nature of DOSI, the effect of smoking appeared to be reversible: while the age- and sex-adjusted DOSI means were higher in current smokers compared to non-smokers, they were indistinguishable between groups of individuals who never smoked and who quit smoking (c.f. [15, 34]).

### Physiological state fluctuations and loss of resilience

To understand the dynamic properties of the organism state fluctuations in relation to aging and diseases, we used two large longitudinal datasets, jointly referred to and available as GEROLONG, including anonymized information on: a) CBC measurements from InVitro, the major Russian clinical diagnostics laboratory and b) physical activity records measured by step counts collected by means of a freely available iPhone application. The CBC slice of the combined dataset included blood test results from 1758 male and 3268 female subjects aged 40 − 90 with complete CBC analyses that were sampled 16 − 20 times within a period of more than three years (up to 42 months).

There was no medical condition information available for the GEROLONG subjects. Hence, for the CBC measurements we used the mean DOSI level corresponding to the “most frail” NHANES and UKB participants as the cutoff value to select “non-frail” GEROLONG individuals (920 male and 1865 female subjects aged 40 − 90) for subsequent analysis.

The difference between the mean DOSI levels in groups of the middle-aged and the eldest available individuals was of the same order as the variation of DOSI across the population at any given age (see Fig. 1B). Accordingly, serial CBC measurements along the individual aging trajectories revealed large stochastic fluctuations of the physiological variables around its mean values, which were considerably different among individual study participants. Naturally, physiological variables at any given moment of time reflect a large number of stochastic factors, such as manifestation of the organism responses to endogenous and external factors (as in Fig. 2C). We therefore focused on the statistical properties of the organism state fluctuations.

Auto-correlation function is the single most important statistical property of a stationary stochastic process represented by a time series *x*(*t*):

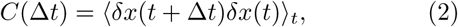

where Δ*t* is the time lag between the subsequent measurements of *x, δx*(*t*) = *x*(*t*) − ⟨*x*⟩_*t*_ is the deviation of *x* from its mean value produced by the averaging ⟨*x*(*t*)⟩_*t*_ along the individual trajectory(see e.g., [35]).

The autocorrelation function of *x* = DOSI averaged over individual trajectories in subsequent age cohorts of GEROLONG dataset was plotted vs. the delay time in Fig. 3A and exhibited exponential decay over a time scale of approximately 2 − 8 weeks depending on age.

**FIG. 3:**
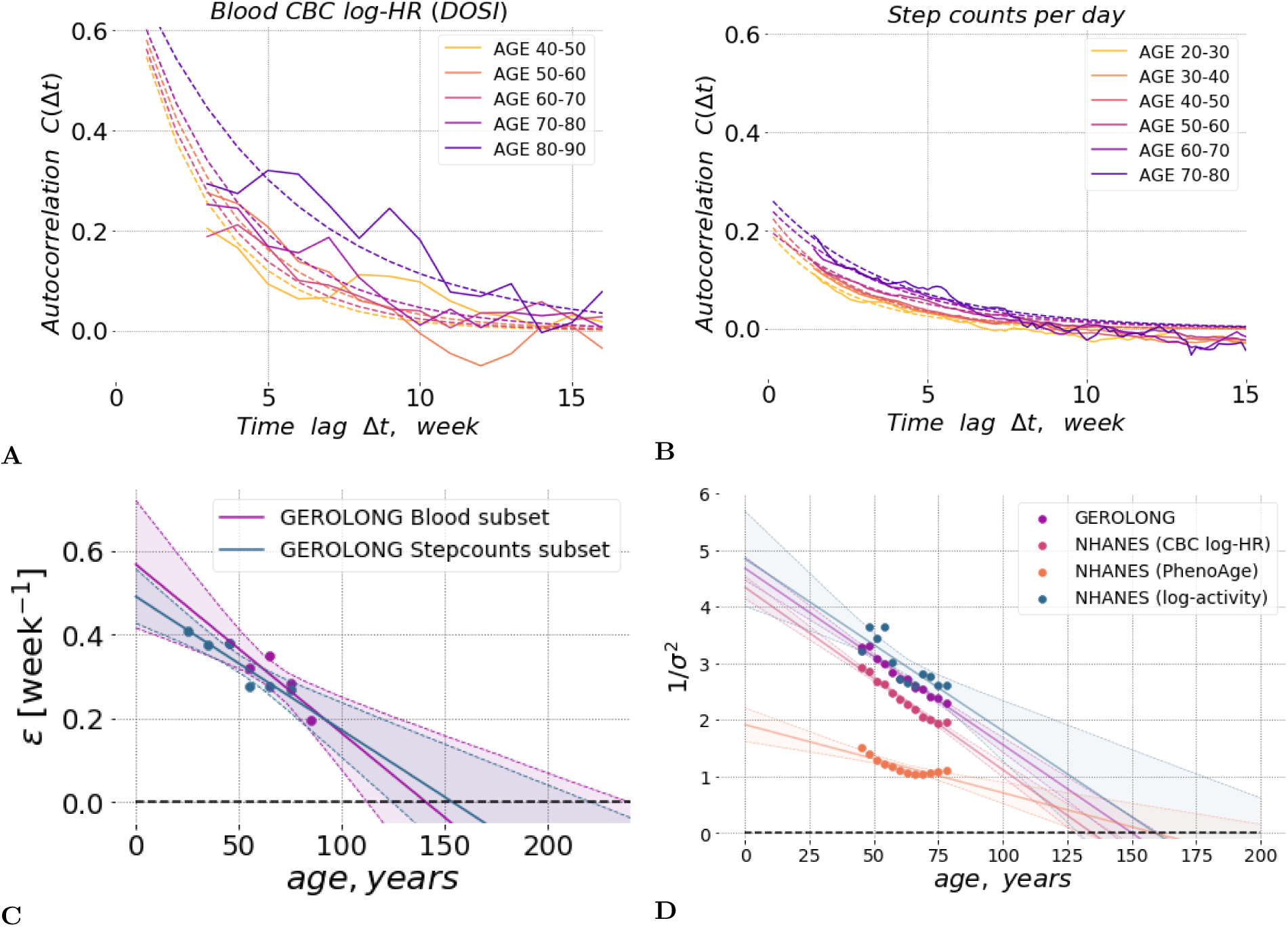
**A**. The auto-correlation function *C*(Δ*t*) of the DOSI fluctuations during several weeks averaged in sequential 10-year age-cohorts of GEROLONG subjects showed gradual age-related remodelling. Experimental data and fit to autocorrelation function are shown with solid and dashed lines, respectively (see details in Supplementary Information). The DOSI correlations are lost over time Δ*t* between the measurements and, hence, the DOSI deviations from its age norm reach the equilibrium distribution faster in younger individuals. **B**. The auto-correlation function *C*(Δ*t*) of fluctuations of the negative logaritm of steps-per-day during several weeks averaged in sequential 10-year age-cohorts of GEROLONG Stepcounts subset subjects showed similar gradual age-related remodelling. **C**. The DOSI relaxation rate (or the inverse characteristic recovery time) computed for sequential age-matched cohorts from the GEROLONG dataset decreased approximately linearly with age and could be extrapolated to zero at an age in the range of *∼*110 − 170 y.o. (at this point, there is complete loss of resilience and, hence, loss of stability of the organism state). The shaded area shows the 95% confidence interval. **D**. The inverse variance of DOSI decreased linearly in all investigated datasets and its extrapolated value vanished (hence, the variance diverged) at an age in the range of 120 − 150 y.o. We performed the linear fit for subjects 40 y.o. and older, excluding the “most frail” (CMI *>* 0.6) individuals. The shaded areas correspond to the 95% confidence intervals. The blue dots and lines show the inverse variance of log-scaled measure of total physical activity (the number of steps per day recorded by a wearable accelerometer) for NHANES participants. Phenoage [25], calculated using explicit age and additional blood biochemistry parameters also demonstrated age-related decrease of the inverse variance in NHANES population.

The exponential character of the autocorrelation function, *C*(Δ*t*) *∼* exp(−*ε*Δ*t*) is a signature of stochastic processes following a simple Langevin equation:

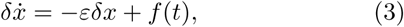

where 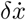 stands for the rate of change in fluctuations *δx, ε* is the relaxation or recovery rate, and *f* is the “force” responsible for deviation of the organism state from its equilibrium.

The auto-correlation function decay time (or simply the auto-correlation time) is inversely proportional to the relaxation (recovery) rate *ε* and characterizes the time scale involved in the equilibration of a system’s state in response to external perturbations (see Supplementary Information). We therefore propose using this quantity as a measure of an organism’s “resilience”, the capacity of an individual organism to resist and recover from the effects of physiological or pathological stresses [36, 37]).

We fitted the DOSI auto-correlation functions averaged over individuals representing subsequent age-matched cohorts to an exponential function of the time delay. We observed that recovery rates obtained from fitting to data in the subsequent age-cohorts decreased approximately linearly with age (Fig. 3C). Extrapolation to older ages suggested that the equilibration rate and hence the resilience is gradually lost over time and is expected to vanish (and hence the recovery time to diverge), at some age of approximately 120 − 150 y.o.).

The exponential decay of auto-correlation function is not merely a peculiarity of an organism state indicator computed from CBC. We were able to use another set of high resolution longitudinal measurements of daily step counts collected by wearable devices. Step counts measurements were obtained from users of fitness wristband (3032 females, 1783 males of age 20 − 85 y.o.). The number of measurements for each user was at least 30 days and up to 5 years.

In [15] we observed, that the variability of physical activity (namely, the logarithm of the average physical activity), that is another hallmark of aging and is associated with age and risks of death or major deceases, also increases with age and hence maybe used as an organism state indicator. The autocorrelation function of the physical activity levels shows already familiar exponential profile and signs of the loss of resilience in subsequent age-matched cohorts as shown in Fig. 3B.

The recovery rate inferred from as the inverse autocorrelation time from physical activity levels trajectories is plotted alongside the recovery rates from CBC-derived DOSI in Fig. 3C. We observed that the recovery rates revealed by the organism state fluctuations measured in apparently unrelated subsystems of the organism (the blood cell counts and physical activity levels) are highly concordant, exhibit the same age-related dynamics.

Eq. (3) predicts, that the variance of DOSI should also increase with age. Indeed, according to the solution of the Langevin equation with a purely random and uncorrelated force, ⟨*f* (*t* +Δ*t*)*f* (*t*)⟩_*t*_ = *Bδ*(Δ*t*) (with *δ*(*x*) being the Dirac’s delta-fucntion and *B* being the power of the stochastic noise), the fluctuations of *x* = DOSI should increase with age thus reflecting the dynamics of the recovery rate: *σ*^2^ ≡ ⟨*δx*^2^⟩ ∼ *B/ϵ*.

Remarkably, the variability in a DOSI did increase with age in every dataset evaluated in this study. Following our theoretical expectations of the inverse relation between the resilience and the fluctuations, we plotted the inverse variance of the DOSI computed in sex- and age-matched cohorts representing the most healthy subjects (see Fig. 3D). Again, extrapolation suggested that, if the tendency holds at older ages, the population variability would increase indefinitely at an age of approximately 120 − 150 y.o.

As expected, the amplification of the fluctuations of the organism state variables with age is not limited to CBC features. In Fig. 3D we plotted the inverse variance of this physical activity feature and found that it linearly decreases with age in such a way that the corresponding variance diverges at the same critical point at the age of approximately 120 − 150 y.o.

To demonstrate the universality of of the organism state dynamics, we followed the fluctuation properties of the Phenoage, another log-linear mortality predictor trained using the explicit age, sex and a number of bio-chemical blood markers [25]. By its nature, PhenoAge is another DOSI produced from a different set of features. Unfortunately, we could not not obtain a sufficient number of individuals with all the relevant markers measurements from the longitudinal dataset from InVitro. Accordingly, we could not compute the corresponding autocorrelation function. We were, however, able to compute PhenoAge for NHANES subjects and observed an increase in variability of the PhenoAge estimate as a function of chronological age and a possible divergence of PhenoAge fluctuations at around the age of 150 y.o.

## DISCUSSION

The simultaneous divergence of the organism state recovery times (critical slowing down in Fig. 3C) and the increasing dynamic range of the the organism state fluctuations (critical fluctuations in Fig. 3D) observed independently in two biological signals is characteristic of proximity of a critical point [23, 35] at some advanced age over 100 y.o. Under these circumstances, the organism state dynamics are stochastic and dominated by the variation of the single dynamic variable (also known as the order parameter) associated with criticality [23, 38].

A proper identification of such a feature requires massive high quality longitudinal measurements and sophisticated approaches auto-regressive models. In a similar study involving CBC variables of aging mice, we were able to obtain an accurate predictor associated with the age, risks of death (and the remaining lifespan), and frailty [39]. In this work we turned the reasoning around and choose to quantify the organism state by the log-linear proportional hazards estimate of the mortality rate followed [15, 25, 40], using CBC and physical activity variables. This inherently dynamic quantitative organism state indicator (DOSI) increased with age, predicted the prospective incidence of age-related diseases and death, and was elevated in cohorts representing typical life-shortening lifestyles, such as smoking, or exhibiting multiple morbidity.

The log-linear risks model predictor demonstrated a non-trivial dependence on age also early in life, that is in the age range with almost no recorded mortality events in the training dataset. The age-cohort averaged DOSI increased and then reached a plateau (Fig. 1B) in good quantitatively consistent with the predictions of the universal theory of ontogenetic growth [27]. According to the theory, the development of any organism is the result of a competition between the production of new tissue and life-sustaining activities. The total amount of the energy available scales as the fractional power of the body mass *m*^*3/4*^ according to the universal allometric Kleiber-West law [41, 42]. On the one hand, the energy requirements for the organism maintenance increase linearly as the body mass grows and hence the initial excess metabolic power drives the growth of the organism until it reaches the dynamic equilibrium corresponding to the mature animal state.

As we can see in Fig. 1B, the mature human organism is dynamically unstable in the long run and deviations from the ontogenetic growth theory predictions pick up slowly well after the organism is fully formed. The organism state dynamics measured by DOSI over lifetime qualitatively reveals at least three regimes reflecting growth, maturation, and aging, respectively. The apparent life-stages correspond well to the results of multi-variate PCA of CBC variance (Fig. 1A) in this work and also that of physical activity acceleration/deceleration patterns from [15]. Every arm of the aging trajectory is characterized by a specific set of features strongly associated with age in the signal.

Schematically, the reported features of the longitudinal organism state dynamics can be summarized with the help of the following qualitative picture (Fig. 4). Far from the critical point (at younger ages), the organism state perturbations can be thought of as confined to the vicinity of a possible stable equilibrium state in a potential energy basin (*A*). Initially, the dynamic stability is provided by a sufficiently high potential energy barrier (*B*) separating this stability basin from the inevitably present dynamically unstable regions (*C*) in the space of physiological parameters. While in stability basin, an organism state experiences stochastic deviation from the meta-stable equilibrium state, which is gradually displaced (see the dashed line *D*) in the course of aging even for the successfully aging individuals.

**FIG. 4:**
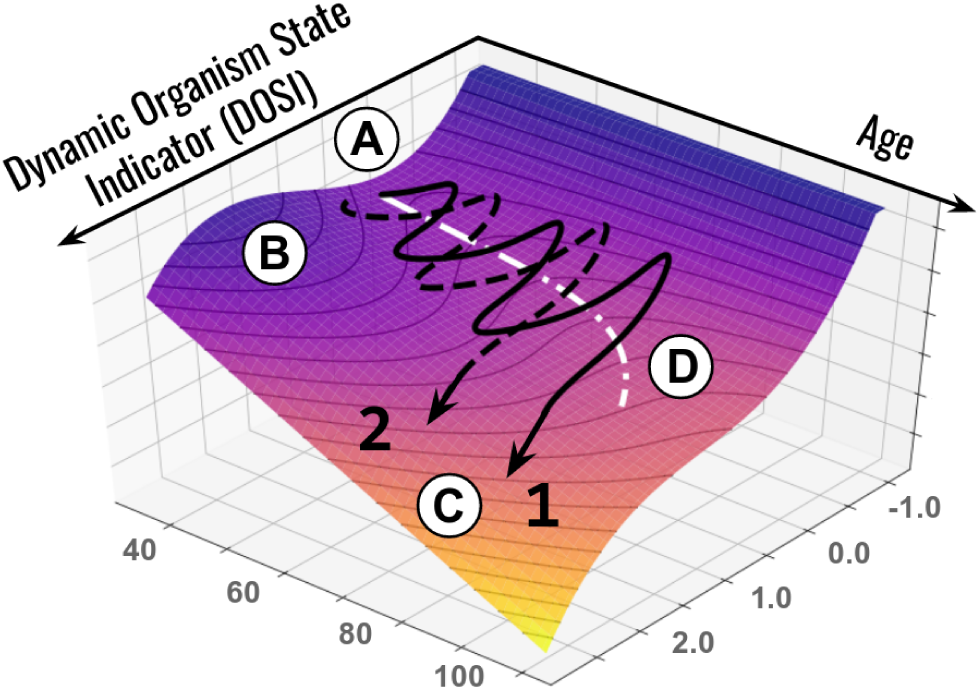
Representative aging trajectories are superimposed over the potential energy landscape (vertical axis) representing regulatory constraints. The stability basin “A” is separated from the unstable region “C” by the potential energy barrier “B”. Aging leads to a gradual decrease in the activation energy and barrier curvature and an exponential increase in the probability of barrier crossing. The stochastic activation into a dynamically unstable (frail) state is associated with acquisition of multiple morbidities and certain death of an organism. The white dotted line “D” represents the trajectory of the attraction basin minimum. Examples 1 (black solid line) and 2 (black dashed line) represent individual life-long stochastic DOSI trajectories that differ with respect to the age of first chronic disease diagnosis.

The characteristic organism state auto-correlation time demonstrated here (3-6 weeks, see Fig. 3A) is much shorter than lifespan. The dramatic separation of time scales makes it very unlikely that the linear decline of the recovery force measured by the recovery rate in Fig. 3C can be explained by the dynamics of the organism state captured by the DOSI variation alone. Therefore, we conclude that the progressive remodeling of the attraction basin geometry reflects adjustment of the DOSI fluctuations to the slow independent process that is aging itself. In this view, the aging drift of the DOSI mean in cohorts of healthy individuals (as in Fig. 1B) is the adaptive organism-level response reflecting, on average, the increasing stress produced by the aging process.

The longitudinal analysis in this work demonstrated that the organism state measured by DOSI follows a stochastic trajectory driven mainly by the organism responses to unpredictable stress factors. Over lifetime, DOSI increases slowsly, on average. The dynamic range of the organism state fluctuations is proportional to the power of noise and is inversely proportional to the recovery rate of the DOSI fluctuations. Therefore, the organism state of healthy individuals at any given age is described by the mean DOSI level, the DOSI variability and its auto-correlation time. Together, the three quantities comprise the minimum set of biomarkers of stress and aging in humans and could be determined and altered, in principle, by different biological mechanisms and therapeutic modalities.

The DOSI recovery rate characterizes fluctuations of DOSI on time scales from few weeks to few months, decreases with age and thus indicates the progressive loss of physiological resilience. Such age-related remodeling of recovery rates has been previously observed in studies of various physiological and functional parameters in humans and other mammals. For example, in humans, a gradual increase in recovery time required after macular surgery was reported in sequential 10-year age cohorts [43] and age was shown to be a significant factor for twelve months recovery and the duration of hospitalization after hip fracture surgery [44, 45], coronary artery bypass [46], acute lateral ankle ligament sprain [47]. A mouse model suggested that the rate of healing of skin wounds also can be a predictor of longevity [48].

The resilience can only be measured directly from high-quality longitudinal physiological data. Framingham Heart Study [7], Dunedin Multidisciplinary Health and Development Study [49] and other efforts produced a growing number of reports involving statistical analysis of repeated measurements from the same persons, see, e.g., [50, 51]. Most of the time, however, the subsequent samples are years apart and hence time between the measurements greatly exceeds the organism state autocorrelation time reported here. This is why, to the best of our understanding, the relation of the organism state recovery rate and mortality has remained largely elusive.

In the presence of stresses, the loss of resilience should lead to destabilization of the organism state. Indeed, in a reasonably smooth potential energy landscape forming the basin of attraction, the activation energy required for crossing the protective barrier (*B*) decreases along with the curvature at the same pace, that is, linearly with age. Whenever the protective barrier is crossed, dynamic stability is lost (see example trajectories 1 and 2 in Fig. 4, which differ by the age of crossing) and deviations in the physiological parameters develop beyond control, leading to multiple morbidities, and, eventually, death.

On a population level, activation into such a frail state is driven by stochastic forces and occurs approximately at the age corresponding to the end of healthspan, understood as “disease-free survival”. Since the probability of barrier crossing is an exponential function of the required activation energy (i.e., the barrier height) [35], the weak coupling between DOSI fluctuations and aging is then the dynamic origin of exponential mortality acceleration known as the Gompertz law. Since the remaining lifespan of an individual in the frail state is short, the proportion of frail subjects at any given age is proportional to the barrier crossing rate, which is an exponential function of age (see Fig. 2B).

The end of healthspan can therefore be viewed as a form of a nucleation transition [35], corresponding in our case to the spontaneous formation of states of chronic diseases out of the metastable phase (healthy organisms). The DOSI is then the order parameter associated with the organism-level stress responses at younger ages and plays the role of the “reaction coordinate” of the transition to the frail state later in life. All chronic diseases and death in our model originate from the dynamic instability associated with single protective barrier crossings. This is, of course, a simplification and yet the assumption could naturally explain why mortality and the incidence of major age-related diseases increase exponentially with age at approximately the same rate [3].

The reduction of slow organism state dynamics to that of a single variable is typical for the proximity of a tipping or critical point [23]. DOSI is therefore the property of the organism as a whole, rather than a characteristics of any specific functional subsystem or organism compartment. We did observe a neat concordance between the organism state recovery rates (Fig. 3C) and DOSI variance divergence (Fig. 3D) from seemingly unrelated sources such as blood markers and the physical activity variables. This is likely a manifestation of common dynamic origin of a substantial part of fluctuations in diverse biological signals ranging from blood markers (CBC and PhenoAge covariates) to physical activity levels.

According to the presented model, external stresses (such as smoking) or diseases produce perturbations that modify the shape of the effective potential leading to the shift of the equilibrium DOSI position. For example, the mean DOSI values in cohorts of individuals who never smoked or who quit smoking are indistinguishable from each other, yet significantly different from (lower than) the mean DOSI in the cohort of smokers (Fig. 2C). Thus, the effect of the external stress factor is reflected by a change in the DOSI and is reversed as soon as the factor is removed.

These findings agree with earlier observations suggesting that the effects of smoking on remaining lifespan and on the risks of developing diseases are mostly reversible once smoking is ceased well before the onset of chronic diseases [15, 34]. The decline in the lung cancer risk after smoking ablation [52] is slower than the recovery rate reported here. This may be the evidence suggesting that long-time stresses may cause hard-to repair damage to the specific tissues and thus produce lasting effects on the resilience.

In the absence of chronic diseases when the organism state is dynamically stable, the elevation of physiological variables associated with the DOSI indicates reversible activation of the most generic protective stress responses. Moderately elevated DOSI levels are therefore protective responses that can measured by molecular markers (e.g., C-reactive protein) and affects general physical and mental health status [40]. On the other hand, the excessive DOSI levels observed in older individuals can be thought of as an aberrant activation of stress-responses beyond the dynamic stability range. This is a characteristics of chronic diseases and death.

We propose that therapies targeting frailty-associated phenotypes (e.g., inflammation) would, therefore, produce distinctly different effects in disease-free versus frail populations. In healthy subjects, who reside in the region of the stability basin (*B*) (see Fig. 4), a treatment-induced reduction of DOSI would quickly saturate over the characteristic auto-correlation time and lead to a moderate decrease in long-term risk of morbidity and death without a change in resilience. Technically, this would translate into an increase in healthspan, although the reduction of health risks would be transient and disappear after cessation of the treatment. In frail individuals, however, the intervention could produce lasting effects and reduce frailty, thus increasing lifespan beyond healthspan. This argument may be supported by longitudinal studies in mice suggesting that the organism state is dynamically unstable, the organism state fluctuations get amplified exponentially at a rate compatible with the mortality rate doubling time, and the effects of transient treatments with life-extending drugs such as rapamycin produce a lasting attenuation of frailty index [39].

The emergence of chronic diseases out of increasingly unstable fluctuations of the organism state provides the necessary dynamic argument to support the derivation of the Gompertz mortality law in the Strehler-Mildvan theory of aging [53]. In [54, 55], the authors suggested that the exponential growth of disease burden observed in the National Population Health Survey of Canadians over 20 y.o. could be explained by an age-related decrease in organism recovery in the face of a constant rate of exposure to environmental stresses. Our study provides evidence suggesting that vanishing resilience cannot be avoided even in the most successfully aging individuals and, therefore, could explain the very high mortality seen in cohorts of super-centennials characterized by the so-called compression of morbidity (late onset of age-related diseases [56]). Formally, such a state of “zero-resilience” at the critical point corresponds to the absolute zero on the vitality scale in the Strehler-Mildvan theory of aging, thus representing a natural limit on human lifespan

The semi-quantitative description of human aging and morbidity proposed here belongs to a class of phenomenological models. Whereas it is possible to associate the variation of the organism state measured by DOSI with the effects of stresses or diseases, the data analysis presented here does not provide any mechanistic explanations for the progressive loss of resilience. It is worth to note that the recent study predicts the maximum human lifespan limit from telomere shortening [57] that is compatible with the estimations presented here. It would therefore be interesting to see if the resilience loss in human cohorts is associated or even caused by the loss of regenerative capacity due to Hayflick limit.

The proximity of the critical point revealed in this work and corresponding to vanishing resilience indicates that the apparent human lifespan limit is not likely to be improved by therapies aimed against specific chronic diseases or frailty syndrome. Thus, no dramatic improvement of the maximum lifespan and hence strong life extension is possible by preventing or curing diseases without interception of the aging process, the root cause of the underlying loss of resilience. We do not foresee any laws of nature prohibiting such an intervention. Therefore, further development of the aging model presented in this work may be a step towards experimental demonstration of a dramatic life-extending therapy.

## MATERIALS AND METHODS

Full details for the materials and methods used in this study, including information of the CBC parameters, Cox proportional hazards model, health conditions, and analysis of physiological state fluctuations are provided in the Supplementary Information.

## SUPPLEMENTARY INFORMATION

### Complete Blood Count Datasets

NHANES CBC data were retrieved from the category “Complete Blood Count with 5-part Differential - Whole Blood” of Laboratory data for NHANES surveys 1999 − 2014. Corresponding UKB CBC data fields with related database codes are listed in Table S1. Samples with missing (or filled with zero) data for any of the used CBC components were discarded. Differential white blood cell percentages were converted to cell counts by multiplication by 0.01 *×* White blood ceel count. All CBC parameters were log-transformed and normalized to zero-mean and unit-variance based on data of NHANES participants aged 40 y.o. and older to further carry out PCA and train Cox proportional hazards model.

### Step Counts Datasets

NHANES step counts per minute records during one week were retrieved from the category “Physical Activity Monitor” of Examination data for NHANES 2005 − 2006 survey.

### Hazards model

The Cox proportional hazards model was trained using NHANES 2015 Public-Use Linked Mortality data. CBC data and mortality linked follow-up available for 40592 NHANES participants aged 18 − 85 y.o. was used. Cox model was trained based on data of participants aged 40 − 85 y.o. (11731 male and 12076 female) with 3792 recorded death events during follow-up until the year 2015 (1999 − 2014 surveys). CBC components and the biological sex label were used as covariates. The model was well-predictive of all-cause mortality and yielded a concordance index value of CI = 0.68 in NHANES and CI = 0.66 in UKB (samples collected 2007 − 2011, 216250 male and 255223 female participants aged 39 − 75 y.o., 13162 recorded death events during follow-up until the year 2016). The Cox proportional hazards model was used as implemented in lifelines package (version 0.14.6) in python. The model was then applied to calculate the hazards ratio for all samples in the GEROLONG, UKB and NHANES cohorts (including individuals younger than 40 y.o.).

The dynamic organism state indicator (DOSI defined as log-hazard ratio of the risk model throughout the manuscript) turned out to be equally well associated with mortality in the NHANES study (HR = 1.43) used for training of the risk model and in the independent UKB study (HR = 1.35; Table S2), which was used as a validation dataset.

### The most prevalent chronic diseases and health status

We quantified the health status of individuals using the sum of major age-related medical conditions that they were diagnosed with, which we termed the compound morbidity index, CMI. The CMI is similar in spirit to the frailty index suggested for NHANES [29]. We were not able to use the frailty index because it was based on Questionnaire and Examination data that were not consistent between all NHANES surveys. Also, we did not have enough corresponding data for the UKB dataset. For CMI determination, we followed [56] and selected the top 11 morbidities strongly associated with age after the age of 40. The list of health conditions included cancer (any kind), cardiovascular conditions (angina pectoris, coronary heart disease, heart attack, heart failure, stroke, or hypertension), diabetes, arthritis and emphysema. Notably, we did not include dementia in the list of diseases since it occurs late in life and hence is severely under-represented in the UKB cohort due to its limited age range. We categorized participants who had more than 6 of those conditions as the “most frail” (CMI *>* 0.6), and those with CMI *<* 0.1 as the “non-frail”. NHANES data for diagnosis with a health condition and age at diagnosis is available in the questionnaire category “Medical Conditions” (MCQ). Data on diabetes and hypertension was retrieved additionally from questionnaire categories “Diabetes” (DIQ) and “Blood Pressure & Cholesterol” (BPQ), respectively.

UK Biobank does not provide aggregated data on these medical conditions. Rather, it provides self-reported questionnaire data (UKB, Category 100074) and diagnoses made during hospital in-patient stay according to ICD10 codes (UKB, Category 2002). We aggregated self-reported and ICD10 (block level) data to match that of NHANES for transferability of the results between populations and datasets. We used the following ICD10 codes to cover the health conditions in UK Biobank: hypertension (I10-I15), arthritis (M00-M25), cancer (C00-C99), diabetes (E10-E14), coronary heart disease (I20-I25), myocardial infarction (I21, I22), angina pectoris (I20), stroke (I60-I64), emphysema (J43, J44), and congestive heart failure (I50).

Consistently with our previous observations in the NHANES and UKB cohorts, DOSI also increased with age in the longitudinal GEROLONG cohort. The average DOSI level as well as its population variance at any given age were, however, considerably larger than those in the reference “non-frail” groups from the NHANES and UKB studies (see Fig. S1A). This difference likely reflects an enrollment bias: many of the GEROLONG blood samples were obtained from patients visiting clinic centers, presumably due to health issues. This could explain why the GEROLONG population appeared generally more frail in terms of DOSI than the reference cohorts of the same age from other studies (Fig. S1A, compare the relative positions of the solid blue line and the two dashed lines representing the GEROLONG cohort and the frail cohorts of the NHANES and UKB studies, respectively).

**TABLE S1:**
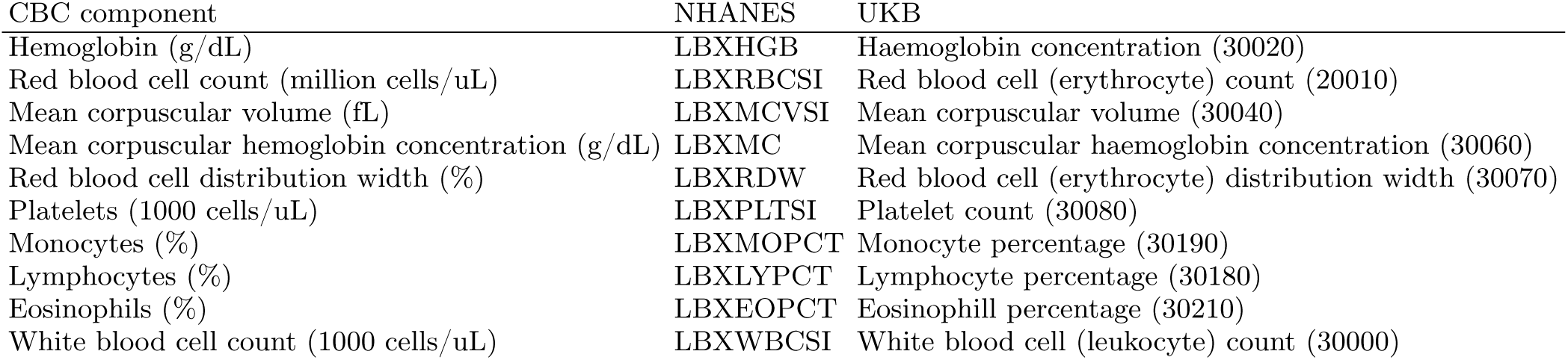
CBC data used in the study.

| CBC component | NHANES | UKB |
| --- | --- | --- |
| Hemoglobin (g/dL) | LBXHGB | Haemoglobin concentration (30020) |
| Red blood cell count (million cells/uL) | LBXRBCSI | Red blood cell (erythrocyte) count (20010) |
| Mean corpuscular volume (fL) | LBXMCVSI | Mean corpuscular volume (30040) |
| Mean corpuscular hemoglobin concentration (g/dL) | LBXMC | Mean corpuscular haemoglobin concentration (30060) |
| Red blood cell distribution width (%) | LBXRDW | Red blood cell (erythrocyte) distribution width (30070) |
| Platelets (1000 cells/uL) | LBXPLTSI | Platelet count (30080) |
| Monocytes (%) | LBXMOPCT | Monocyte percentage (30190) |
| Lymphocytes (%) | LBXLYPCT | Lymphocyte percentage (30180) |
| Eosinophils (%) | LBXEOPCT | Eosinophill percentage (30210) |
| White blood cell count (1000 cells/uL) | LBXWBCSI | White blood cell (leukocyte) count (30000) |

**TABLE S2:**
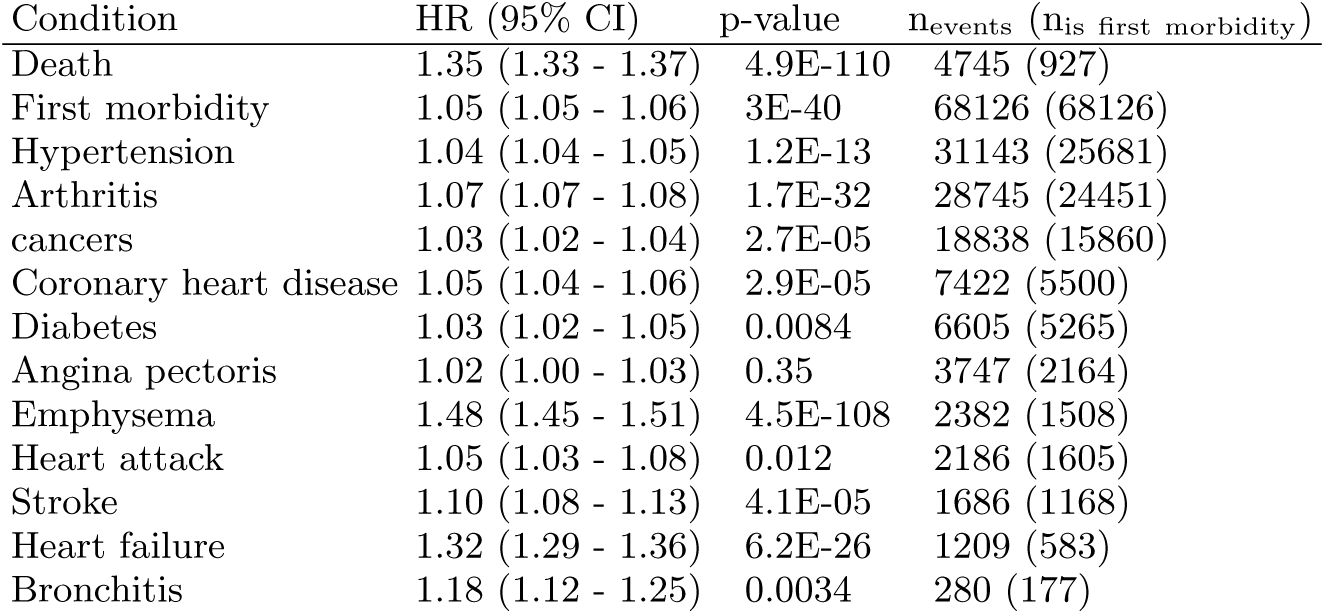
Significance of prediction of acquiring a health condition based on estimated log hazards ratio (adjusted for age and gender). Only UKB subjects with none of the listed health conditions at the time of survey were considered; the total number of subjects evaluated for each condition was 263956. The numbers in parentheses in the far right column indicate the occurrence of the disease being the first diagnosis in an individual.

| Condition | HR (95% CI) | p-value | n<sub>events</sub> (n<sub>is first morbidity</sub>) |
| --- | --- | --- | --- |
| Death | 1.35 (1.33 - 1.37) | 4.9E-110 | 4745 (927) |
| First morbidity | 1.05 (1.05 - 1.06) | 3E-40 | 68126 (68126) |
| Hypertension | 1.04 (1.04 - 1.05) | 1.2E-13 | 31143 (25681) |
| Arthritis | 1.07 (1.07 - 1.08) | 1.7E-32 | 28745 (24451) |
| cancers | 1.03 (1.02 - 1.04) | 2.7E-05 | 18838 (15860) |
| Coronary heart disease | 1.05 (1.04 - 1.06) | 2.9E-05 | 7422 (5500) |
| Diabetes | 1.03 (1.02 - 1.05) | 0.0084 | 6605 (5265) |
| Angina pectoris | 1.02 (1.00 - 1.03) | 0.35 | 3747 (2164) |
| Emphysema | 1.48 (1.45 - 1.51) | 4.5E-108 | 2382 (1508) |
| Heart attack | 1.05 (1.03 - 1.08) | 0.012 | 2186 (1605) |
| Stroke | 1.10 (1.08 - 1.13) | 4.1E-05 | 1686 (1168) |
| Heart failure | 1.32 (1.29 - 1.36) | 6.2E-26 | 1209 (583) |
| Bronchitis | 1.18 (1.12 - 1.25) | 0.0034 | 280 (177) |

**FIG. S1:**
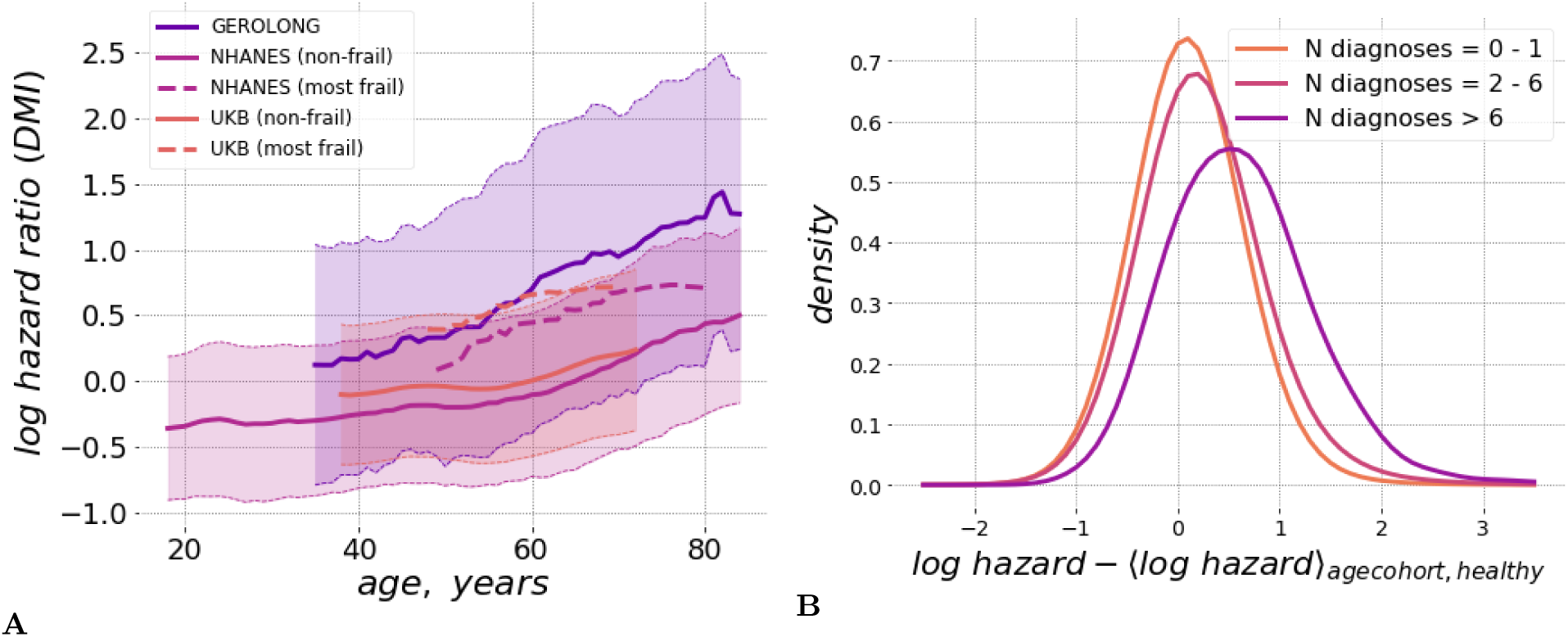
**A**. DOSI mean values (lines) and variance (shaded areas) are plotted relative to age for the NHANES (same as in Fig. 1B), UKB and GEROLONG datasets (color-matched with respect to each study). For NHANES and UKB the solid line and shaded regions mark the population average ad the range spanned by one standard deviation from it for the “non-frail” (CMI *<* 0.1) participats. The population mean for the “most frail” (CMI *>* 0.6) individuals is shown with dashed lines. **B**. Distributions of sex- and age-adjusted DOSI in cohorts of UKB participants in different morbidity categories relative to the DOSI mean in cohorts of “non-frail” (one or no diagnoses, CMI *<* 0.1) individuals. Note that the distribution function in the “most frail” group (more than 6 diagnoses, CMI *>* 0.6) exhibited the largest shift and a profound deviation from the symmetric form, similarly as it was seen in NHANES.

## References

[1] Arnold Mitnitski and Kenneth Rockwood. The rate of aging: the rate of deficit accumulation does not change over the adult life span. Biogerontology, 17(1):199–204, 2016.

[2] Ruby Yu, Wan-Chi Wu, Jason Leung, Susan C Hu, and Jean Woo. Frailty and its contributory factors in older adults: a comparison of two asian regions (hong kong and taiwan). International journal of environmental research and public health, 14(10):1096, 2017.

[3] Aleksandr Zenin, Yakov Tsepilov, Sodbo Sharapov, Evgeny Getmantsev, Leonid Menshikov, Peter Fedichev, and Yurii Aulchenko. Identification of 12 genetic loci associated with human healthspan. bioRxiv, page 300889, 2018.

[4] Dmitriy I Podolskiy, Alexei V Lobanov, Gregory V Kryukov, and Vadim N Gladyshev. Analysis of cancer genomes reveals basic features of human aging and its role in cancer development. Nature communications, 7:12157, 2016.

[5] Teresa Niccoli and Linda Partridge. Ageing as a risk factor for disease. Current Biology, 22(17):R741–R752, 2012.

[6] Nir Barzilai and Gad Rennert. The rationale for delaying aging and the prevention of age-related diseases. Rambam Maimonides medical journal, 3(4), 2012.

[7] Thomas R Dawber, Gilcin F Meadors, and Felix E Moore Jr. Epidemiological approaches to heart disease: the framingham study. American Journal of Public Health and the Nations Health, 41(3):279–286, 1951.

[8] Cathie Sudlow, John Gallacher, Naomi Allen, Valerie Beral, Paul Burton, John Danesh, Paul Downey, Paul Elliott, Jane Green, Martin Landray, et al. Uk biobank: an open access resource for identifying the causes of a wide range of complex diseases of middle and old age. PloS medicine, 12(3):e1001779, 2015.

[9] Paul Brennan, Markus Perola, Gert-Jan van Ommen, Elio Riboli, European Cohort Consortium, et al. Chronic disease research in europe and the need for integrated population cohorts. European journal of epidemiology, 32(9):741–749, 2017.

[10] M. E. Levine. Modeling the rate of senescence: can estimated biological age predict mortality more accurately than chronological age? J. Gerontol. A Biol. Sci. Med. Sci., 68(6):667–674, Jun 2013.

[11] G. Hannum, J. Guinney, L. Zhao, L. Zhang, G. Hughes, S. Sadda, B. Klotzle, M. Bibikova, J. B. Fan, Y. Gao, R. Deconde, M. Chen, I. Rajapakse, S. Friend, T. Ideker, and K. Zhang. Genome-wide methylation profiles reveal quantitative views of human aging rates. Mol. Cell, 49(2):359–367, Jan 2013.

[12] S. Horvath. DNA methylation age of human tissues and cell types. Genome Biol., 14(10):R115, 2013.

[13] Philippe Terrier and Fabienne Reynard. Effect of age on the variability and stability of gait: a cross-sectional treadmill study in healthy individuals between 20 and 69 years of age. Gait & posture, 41(1):170–174, 2015.

[14] Tim Althoff, Jennifer L Hicks, Abby C King, Scott L Delp, Jure Leskovec, et al. Large-scale physical activity data reveal worldwide activity inequality. Nature, 547(7663):336, 2017.

[15] Timothy V. Pyrkov, Evgeny Getmantsev, Boris Zhurov, Konstantin Avchaciov, Mikhail Pyatnitskiy, Leonid Menshikov, Kristina Khodova, Andrei V. Gudkov, and Peter O. Fedichev. Quantitative characterization of biological age and frailty based on locomotor activity records. Aging, 10(10):2973–2990, 2018.

[16] Juulia Jylhävä, Nancy L. Pedersen, and Sara Hägg. Biological Age Predictors. EBioMedicine, 21:29–36, 2017.

[17] Benjamin Gompertz. A sketch of an analysis and notation applicable to the value of life contingencies. Philosophical Transactions of the Royal Society, 110:214–294, 1820.

[18] William Matthew Makeham. On the law of mortality and construction of annuity tables. The Assurance Magazine and Journal of the Institute of Actuaries, 8(06):301–310, 1860.

[19] Carlos López-Otín, Maria A Blasco, Linda Partridge, Manuel Serrano, and Guido Kroemer. The hallmarks of aging. Cell, 153(6):1194–1217, 2013.

[20] Sanne MW Gijzel, Heather E Whitson, Ingrid A van de Leemput, Marten Scheffer, Dieneke van Asselt, Jerrald L Rector, Marcel GM Olde Rikkert, and René JF Melis. Resilience in clinical care: Getting a grip on the recovery potential of older adults. Journal of the American Geriatrics Society, 2019.

[21] Heather E Whitson, Wei Duan-Porter, Kenneth E Schmader, Miriam C Morey, Harvey J Cohen, and Cathleen S Colón-Emeric. Physical resilience in older adults: systematic review and development of an emerging construct. Journals of Gerontology Series A: Biomedical Sciences and Medical Sciences, 71(4):489–495, 2015.

[22] Olde Rikkert, GM Marcel, Vasilis Dakos, Timothy G Buchman, Rob de Boer, Leon Glass, Angélique OJ Cramer, Simon Levin, Egbert van Nes, George Sugihara, et al. Slowing down of recovery as generic risk marker for acute severity transitions in chronic diseases. Critical care medicine, 44(3):601–606, 2016.

[23] Marten Scheffer, Jordi Bascompte, William A Brock, Victor Brovkin, Stephen R Carpenter, Vasilis Dakos, Hermann Held, Egbert H Van Nes, Max Rietkerk, and George Sugihara. Early-warning signals for critical transitions. Nature, 461(7260):53–59, 2009.

[24] Marten Scheffer. Complex systems: foreseeing tipping points. Nature, 467(7314):411, 2010.

[25] Zuyun Liu, Pei-Lun Kuo, Steve Horvath, Eileen Crimmins, Luigi Ferrucci, and Morgan Levine. Phenotypic age: a novel signature of mortality and morbidity risk. bioRxiv, page 363291, 2018.

[26] Timothy V Pyrkov, Konstantin Slipensky, Mikhail Barg, Alexey Kondrashin, Boris Zhurov, Alexander Zenin, Mikhail Pyatnitskiy, Leonid Menshikov, Sergei Markov, and Peter O Fedichev. Extracting biological age from biomedical data via deep learning: too much of a good thing? Scientific reports, 8(1):5210, 2018.

[27] Geoffrey B West, James H Brown, and Brian J Enquist. A general model for ontogenetic growth. Nature, 413(6856):628–631, 2001.

[28] David R Cox. Regression models and life-tables. In Breakthroughs in statistics, pages 527–541. Springer, 1992.

[29] K Rockwood, JM Blodgett, O Theou, MH Sun, HA Feridooni, A Mitnitski, RA Rose, J Godin, E Gregson, and SE Howlett. A frailty index based on deficit accumulation quantifies mortality risk in humans and in mice. Scientific Reports, 7, 2017.

[30] Benjamin Gompertz. On the nature of the function expressive of the law of human mortality, and on a new mode of determining the value of life contingencies. Philosophical transactions of the Royal Society of London, 115:513–583, 1825.

[31] Andrea Ganna and Erik Ingelsson. 5 year mortality predictors in 498 103 uk biobank participants: a prospective population-based study. The Lancet, 386(9993):533–540, 2015.

[32] R OâĂŹdonnell, D Breen, S Wilson, and R Djukanovic. Inflammatory cells in the airways in copd. Thorax, 61(5):448–454, 2006.

[33] Tariq Bhat, Sumaya Teli, Jharendra Rijal, Hilal Bhat, Muhammad Raza, Georges Khoueiry, Mustafain Meghani, Muhammad Akhtar, and Thomas Costantino. Neutrophil to lymphocyte ratio and cardiovascular diseases: a review. Expert review of cardiovascular therapy, 11(1):55–59, 2013.

[34] Donald H Taylor Jr, Vic Hasselblad, S Jane Henley, Michael J Thun, and Frank A Sloan. Benefits of smoking cessation for longevity. American journal of public health, 92(6):990–996, 2002.

[35] LD Landau and EM Lifshitz. Physical kinetics, vol. 10. Course of Theoretical Physics, 1981.

[36] Gregory Hicks and Ram R Miller. Physiological resilience. In Resilience in Aging, pages 89–103. Springer, 2011.

[37] N Jennifer Klinedinst and Alisha Hackney. Physiological resilience and the impact on health. In Resilience in Aging, pages 105–131. Springer, 2018.

[38] Baruch Barzel and Albert-László Barabási. Universality in network dynamics. Nature physics, 9(10):673–681, 2013.

[39] Konstantin Avchaciov, Marina P Antoch, Ekaterina L Andrianova, Andrei E Tarkhov, Leonid I Menshikov, Andrei V Gudkov, and Peter O Fedichev. Identification of a blood test-based biomarker of aging through deep learning of aging trajectories in large phenotypic datasets of mice. bioRxiv, 2020.

[40] Timothy V Pyrkov and Peter O Fedichev. Biological age is a universal marker of aging, stress, and frailty. bioRxiv, page 578245, 2019.

[41] Max Kleiber et al. Body size and metabolism. Hilgardia, 6(11):315–353, 1932.

[42] Geoffrey B West, James H Brown, and Brian J Enquist. A general model for the origin of allometric scaling laws in biology. Science, 276(5309):122–126, 1997.

[43] Yonguk Kim, Eung Suk Kim, Seung-Young Yu, and Hyung Woo Kwak. Age-related clinical outcome after macular hole surgery. Retina, 37(1):80–87, 2017.

[44] Jana M Mossey, Elizabeth Mutran, Kathryn Knott, and Rebecca Craik. Determinants of recovery 12 months after hip fracture: the importance of psychosocial factors. American journal of public health, 79(3):279–286, 1989.

[45] Kenneth J Koval, Mary Louise Skovron, Gina B Aharonoff, and Joseph D Zuckerman. Predictors of functional recovery after hip fracture in the elderly. Clinical orthopaedics and related research, 1(348):22–28, 1998.

[46] NT Artinian, C Duggan, and P Miller. Age differences in patient recovery patterns following coronary artery bypass surgery. American Journal of Critical Care, 2(6):453–461, 1993.

[47] Jacqueline Yewande Thompson, Christopher Byrne, Mark A Williams, David J Keene, Micheal Maia Schlus- sel, and Sarah E Lamb. Prognostic factors for recovery following acute lateral ankle ligament sprain: a systematic review. BMC musculoskeletal disorders, 18(1):421, 2017.

[48] Hagai Yanai, Arie Budovsky, Robi Tacutu, and Vadim E Fraifeld. Is rate of skin wound healing associated with aging or longevity phenotype? Biogerontology, 12(6):591–597, 2011.

[49] Daniel W Belsky, Avshalom Caspi, Renate Houts, Harvey J Cohen, David L Corcoran, Andrea Danese, HonaLee Harrington, Salomon Israel, Morgan E Levine, Jonathan D Schaefer, et al. Quantification of biological aging in young adults. Proceedings of the National Academy of Sciences, 112(30):E4104–E4110, 2015.

[50] Ayelet Alpert, Yishai Pickman, Michael Leipold, Yael Rosenberg-Hasson, Xuhuai Ji, Renaud Gaujoux, Hadas Rabani, Elina Starosvetsky, Ksenya Kveler, Steven Schaffert, et al. A clinically meaningful metric of immune age derived from high-dimensional longitudinal monitoring. Nature medicine, 25(3):487, 2019.

[51] Sara Ahadi, Wenyu Zhou, Sophia Miryam Schüssler-Fiorenza Rose, M Reza Sailani, Kévin Contrepois, Monika Avina, Melanie Ashland, Anne Brunet, and Michael Snyder. Personal aging markers and ageotypes revealed by deep longitudinal profiling. Nature Medicine, 26(1):83–90, 2020.

[52] Hilary A Tindle, Meredith Stevenson Duncan, Robert A Greevy, Ramachandran S Vasan, Suman Kundu, Pierre P Massion, and Matthew S Freiberg. Lifetime smoking history and risk of lung cancer: Results from the framingham heart study. JNCI: Journal of the National Cancer Institute, 110(11):1201–1207, 2018.

[53] Bernard L Strehler and Albert S Mildvan. General theory of mortality and aging. Science See Saiensu, 132, 1960.

[54] Arnold Mitnitski, Xiaowei Song, and Kenneth Rockwood. Assessing biological aging: the origin of deficit accumulation. Biogerontology, 14(6):709–717, 2013.

[55] Arnold Mitnitski and Kenneth Rockwood. Aging as a process of deficit accumulation: its utility and origin. In Aging and Health-A Systems Biology Perspective, volume 40, pages 85–98. Karger Publishers, 2015.

[56] Stacy L Andersen, Paola Sebastiani, Daniel A Dworkis, Lori Feldman, and Thomas T Perls. Health span approximates life span among many supercentenarians: compression of morbidity at the approximate limit of life span. Journals of Gerontology Series A: Biomedical Sciences and Medical Sciences, 67(4):395–405, 2012.

[57] Kurt Whittemore, Elsa Vera, Eva Martínez-Nevado, Carola Sanpera, and Maria A Blasco. Telomere shortening rate predicts species life span. Proceedings of the National Academy of Sciences, 116(30):15122–15127, 2019.

